# Modeling of astrocyte networks: towards realistic topology and dynamics

**DOI:** 10.1101/2020.12.20.423646

**Authors:** D. V. Verveyko, A. Yu. Verisokin, D. E. Postnov, A. R. Brazhe

**Affiliations:** Department of Theoretical Physics, Kursk State University, Radishcheva st., 33, 305000, Kursk, Russia; Saratov State University, Astrakhanskaya st., 83, 410012, Saratov, Russia; Department of Biophysics, Biological Faculty, Lomonosov Moscow State University, Leninskie Gory, 1/24, 119234 Moscow, Russia; Department of Molecular Neurobiology, Institute of Bioorganic Chemistry RAS, 117997, Russian Federation, Moscow, GSP-7, Ulitsa Miklukho-Maklaya, 16/10

**Author notes:** Correspondence: A. R. Brazhe.

**Keywords:** calcium signaling, cell morphology, noise-driven dynamics

## Abstract

Neuronal firing and neuron-to-neuron synaptic wiring are currently widely described as orchestrated by astrocytes — elaborately ramified glial cells tiling the cortical and hippocampal space into non-overlapping domains, each covering hundreds of individual dendrites and hundreds thousands synapses. A key component to astrocytic signaling is the dynamics of cytosolic Ca^2+^ which displays multiscale spatiotemporal patterns from short confined elemental Ca^2+^ events (puffs) to Ca^2+^ waves expanding through many cells. Here we synthesize the current understanding of astrocyte morphology, coupling local synaptic activity to astrocytic Ca^2+^ in perisynaptic astrocytic processes and morphology-defined mechanisms of Ca^2+^ regulation in a distributed model. To this end, we build simplified realistic data-driven spatial network templates and compile model equations as defined by local cell morphology. The input to the model is spatially uncorrelated stochastic synaptic activity. The proposed modeling approach is validated by statistics of simulated Ca^2+^ transients at a single cell level. In multicellular templates we observe regular sequences of cell entrainment in Ca^2+^ waves, as a result of interplay between stochastic input and morphology variability between individual astrocytes. Our approach adds spatial dimension to the existing astrocyte models by employment of realistic morphology while retaining enough flexibility and scalability to be embedded in multiscale heterocellular models of neural tissue. We conclude that the proposed approach provides a useful description of neuron-driven Ca^2+^-activity in the astrocyte syncytium.

## 1 INTRODUCTION

Astrocytes of the cortical and hippocampal gray matter are important actors in a number of information processing processes, including synaptic plasticity, long-term potentiation, and synchronization of neuronal firing (De Pitta et al, 2015; Haydon, 2001; Lee et al, 2014; Poskanzer and Yuste, 2016) as well as in coupling neuronal activity to blood flow changes (Otsu et al, 2015). Recent evidence converges on a close connection of these functions with whole-brain processes and systemic regulation pathways. Thus, astrocytes respond to and are able to regulate systemic blood pressure Marina et al (2020); they significantly (up to 60 %) change their volume during sleep or under anesthesia Xie et al (2013); astrocytes play an important role in the clearance of beta-amyloids, a process with mechanisms that are now being actively discussed Iliff et al (2012); Mestre et al (2020); Abbott et al (2018); Semyachkina-Glushkovskaya et al (2018); both intracellular and network-level activity of astrocytes are significantly different in sleep and during wakefulness, and activates with locomotion Ingiosi et al (2020); Bojarskaite et al (2020); McCauley et al (2020). It is important to note that many of the mentioned astrocyte functions are not directly related to neural activity, but are governed by their own regulatory pathways O’Donnell et al (2015). Some of these functions are tightly linked to dynamic regulation of astrocyte morphology and volume and depend, for example, on the circadian rhythm of aquaporin expression Hablitz et al (2020).

In summary, this frames a new mindset for understanding the function of astrocytes and at the same time poses a challenge for modeling studies. Namely, the morphological features should now be considered as a specific control parameter that significantly contribute to the both single-cell dynamics and network activity patterns. This problem, breaks down into three specific tasks: (i) to provide tractable, but still biologically reasonable mathematical account for contribution of subcellular morphological features to intracellular calcium dynamics; (ii) to move away from the physical “lattice” approach in modeling to employment of irregular structures observed in the experiment, both for an individual cell and for a network; (iii) to reveal how realistic morphological features are manifested in the spatiotemporal patterns of the calcium dynamics.

We address these tasks in more detail in the rest of the Introduction.

### 1.1 Calcium signaling in astrocytes

With plasma membranes enriched in a battery of K^+^-channels and lacking 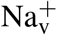, astrocytes are not electrically excitable. On the other hand, they display a rich repertoire of Ca^2+^-activity at multiple spatial and temporal scales (Volterra et al, 2014; Lind et al, 2013; Wu et al, 2014; Bindocci et al, 2017; Stobart et al, 2018). Although astrocytic Ca^2+^ transients can occur spontaneously, their frequency is modulated by neuronal activity, changes in local tissue oxygenation, and other factors. As outputs, Ca^2+^-activity in astrocytes leads to release of signaling molecules: gliotransmitters, such as GABA, D-serine, and glutamate, as well as vasoactive metabolites (Bazargani and Attwell, 2016). Recent experimental evidence obtained with genetically encoded or pipette-loaded Ca^2+^ indicators (Tong et al, 2013; Rungta et al, 2016) heralds functional segregation between the less frequent global internal store-operated Ca^2+^ transients at the level of cell soma and primary branches, and the more frequent spatially limited microdomain Ca^2+^ transients in the thin mesh of astrocytic leaflets — ramified nanoscopic processes, also known as perisynaptic processes (PAPs) due to their proximity to synaptic connections between neurons. The transients located in the leaflets primarily rely on influx of Ca^2+^ through plasma membrane, in part because of the high surface-to-volume ratio in this region and in part because the leaflets are often devoid of organelles including ER (Patrushev et al, 2013) and thus can’t support exchange with intracellular stores.

The coupling from synaptic activity to local Ca^2+^ transients in PAPs and from the latter to global Ca^2+^ events is an area of active research. One plausible causal pathway can be formulated as follows (Rojas et al, 2007; Verkhratsky et al, 2012; Parpura et al, 2016; Kirischuk et al, 2016): neurotransmitters, released from the presynaptic membranes, primarily glutamate, but also GABA, are cleared from the extracellular space by astrocytic transporters utilizing Na^+^ gradient to drive the neurotransmitters into the cell. This leads to build up of Na^+^ ions in the cytosol, which can lead to temporary reversal of Na^+^/Ca^2+^-exchanger allowing for Ca^2+^ entry via this transporter. If this local Ca^2+^ influx happens near the ER and coincides with an increase in inositol trisphosphate production by phospholipase C, it can trigger Ca^2+^-induced release of Ca^2+^ from intracellular stores via IP_3_ receptors (IP_3_Rs) of the ER.

The release of calcium from ER is spatially inhomogeneous due to the non-uniform, clustered, distribution of IP_3_ receptors (Ross, 2012; Smith et al, 2009; Taufiq-Ur-Rahman et al, 2009), with clusters spaced at about 0.5–5 *µ*m apart. At a detailed level, calcium release from the receptor clusters has a stochastic character. The effect of the stochastic activation of IP_3_R clusters on the calcium dynamics has been investigated by Shuai and Jung both in point and distributed models (Shuai and Jung, 2002, 2003). In the case of a large enough number of clusters, Ca^2+^ release events can be averaged to a lumped deterministic description. Particularly, the increase in IP_3_ level transforms stochastic calcium increases into regular waves.

Recapitulating, calcium signaling mechanisms are inhomogeneous across the cell and depend on local morphological parameters, which has to be taken into account in modeling. It seems practical to introduce a metaparameter to describe the relative inputs of store-related and plasma membrane-related Ca^2+^ pathways. This metaparameter can reflect local surface-to-volume ratio or the dominant size of processes and can empirically be linked to the astrocyte cytoplasm volume fraction parameter, which can be estimated directly from fluorescent images.

### 1.2 Cell morphology and network connectivity

Astrocytes have intricate and highly complex morphology, which raises computational issues and demands an elaborate approach to modeling. The contribution of the astrocytic spatial segregation and coupling to brain physiology and functions is still not sufficiently understood, especially taking into account that astrocyte-to-neuron and astrocyte-to-astrocyte interaction mechanisms are diverse and depend on brain region. The existence of intercellular Ca^2+^ waves traveling across the network of astrocytes suggests a distinct mechanism for long-distance signaling (Cornell-Bell et al, 1990) and plasticity, which operates in parallel to and at much slower time scales than neuronal synaptic transmission (Pirttimaki and Parri, 2013; Sims et al, 2015).

The size of cliques of cortical astrocytes coupled within a local network is estimated around 60–80 cells (Houades et al, 2006, 2008; Haas et al, 2006), but several networks can also connect via a limited number of “hub” astrocytes (Carmignoto, 2000). The implications of inter-astrocyte connectivity have been analysed in a modeling study by Lallouette et al (2014) with the main conclusion that sparse short-range connections can promote Ca^2+^ wave propagation along the network. This allows to conjecture that once initiated, a wave of excitation can propagate over long distances in the brain cortex and affect (activate or inhibit) postsynaptic neurons at distant synaptic terminals, although most Ca^2+^ events are confined to a single astrocyte spatial domain. Propagating calcium waves can travel distances of more than 100 *µ*m with speed from 7 to 27 *µ*m/sec in culture and brain slices (Dani et al, 1992). However, the waves observed *in vivo* rarely spread more than 80 *µ*m (Hoogland et al, 2009; Brazhe et al, 2013), although this observation can be influenced by imaging protocol, as Kuga and colleagues reported large-scale Ca^2+^ glissandi *in vivo* that were only observable under low laser intensity (Kuga et al, 2011).

It follows that for meso-scale problems related to brain tissue physiology, it is computationally cumbersome to build a ground-up model starting from individual processes. We propose a more pragmatic approach based on texture-like volume segmentation to classes such as “soma”, “large branches” and “gliapil” or a mesh of unresolved thin processes. This rasterization radically simplifies model implementation and scales to large networks. At the same time, by defining morphology-based spatial distribution of a metaparameter, one can study the effects of spatial heterogeneity at different scales. Indeed, the spatial distributions used for simulations are ideally data-driven. Because it is not always possible to infer the astrocytic network structure or even individual domain boundaries from experimental data, and because the networks can be variable anyway, it seems inviting to generate variable astrocytic tilings from images of individual cells.

### 1.3 Modelling studies

Models of IP_3_-mediated Ca^2+^ oscillations have been extensively reviewed both in general (Dupont et al, 2011) and in application to astrocytes (Riera et al, 2011; Manninen and Havela, 2017; Oschmann et al, 2017a), which included both point- and spatially extended models. In particular, the De Young – Keizer model stemmed several currently popular models of Ca^2+^ dynamics in literature. This model allows to simulate IP_3_-sensitive calcium dynamics in cytoplasm and ER occurring at the constant level of IP_3_ including also a variant of the model with the positive-feedback mechanism of Ca^2+^ on IP_3_ production (De Young and Keizer, 1992). Li and Rinzel (Li and Rinzel, 1994) reduced De Young – Keizer model to a two-variable system introducing the experimentally observed time scale difference between fast and slow inactivation of IP_3_ receptor by Ca^2+^. Adding the dynamics for [IP_3_] with synthesis dependent on activation of metabotropic glutamate receptors and [Ca^2+^] degradation leads to a three variable model (Ullah et al, 2006). Also building on Li – Rinzel model and providing a more detailed description of IP_3_ degradation, De Pitta and co-authors proposed a three-variable model for glutamate-induced intracellular calcium dynamics caused by the synaptic activity in astrocytes (De Pitta et al, 2009). One of the first models for intercellular propagation of calcium waves has been described in (Sneyd et al, 1994) by adding diffusion of IP_3_ and cytosolic Ca^2+^ to the two-pool Ca^2+^ model.

Recently Savtchenko et al (2018) suggested an advanced NEURON-based modeling environment for detailed spatially extended models of astrocytes. However, they didn’t address full calcium dynamics models or morphology-defined variations of mechanisms. Specifically, the relative weights of plasma membrane-dependent mechanisms (IP_3_ synthesis and Ca^2+^ influx) and store-dependent mechanisms scale with astrocytic process morphology, as defined by surface to volume ratio, cytoplasm volume fraction and the physical presence of ER in the process. This has been studied in point-models by Oschmann et al (2017b) and in 1D extended model by Wu et al (2018). Recently, Brazhe et al (2018) studied the implications of the spatial segregation between IP_3_ synthesis and plasma membrane exchange and the IP_3_-mediated ER exchange in discrete spatial templates of variable complexity.

### 1.4 The proposed modeling approach

We present a model of multi-cellular network of astrocytes based on realistic spatial templates. Our single-cell model is considerably simpler than in Savtchenko et al (2018), allowing for smaller computational costs.

Here we focus on the implications of the morphology-dependent spatial segregation of the Ca^2+^ signaling mechanisms between astrocytic leaflets and branches. We thus follow the lines set out in Brazhe et al (2018) towards more realistic and larger scale spatial templates, ranging from single astrocytes to networks. The rest of the paper is organized as follows: we start from a description of the proposed model in a top-down order: the general concept is followed by proposed algorithm of creating spatial templates for modeling and then continues with description of the differential equations for dynamics of intracellular and ER Ca^2+^, intracellular IP_3_, and extracellular glutamate concentrations. Having defined the model, we test its plausibility on single-astrocyte templates and after quantification of Ca^2+^ event statistics we proceed to behavior of astrocyte networks, where we observe noise-driven regular activation patterns.

## 2 MODEL

### 2.1 Model design and overview

In this work we aim to conceptualize our current understanding of spatial organization of the astrocytic Ca^2+^ dynamics in a form of a spatially detailed model of individual and networked astrocytes excited by stochastic background neuronal activity. In the light of the striking differences between Ca^2+^ signaling in astrocytic leaflets and thin processes on the one hand and global somatic signaling on the other, we start with segregation of the modeling space into three major classes as shown in Figure 1: astrocyte soma with thick branches (I), a mesh of astrocytic thin processes (II) and extracellular space (III). The continuum between the two extreme classes I and II is defined as local fraction of astrocytic cytoplasm volume (AVF) and a related parameter — local surface-to-volume ratio (SVR) of the astrocytic processes. In extreme class I regions, such as soma, Ca^2+^ dynamics is dominated by exchange with intracellular stores, and a unit of modeled space (template pixel/voxel) contains only astrocyte, while in extreme class II regions (leaflets), Ca^2+^ dynamics is dominated by exchange with plasma membrane and each modeled pixel contains a mesh of extremely thin astrocyte processes tangled with neuropil. We thus define a mapping of each pixel in the spatial model template to either class III (no astrocyte) or to a continuous variable between the extreme cases of class I and II with implications in local calcium dynamics and diffusion.

**Figure 1.**
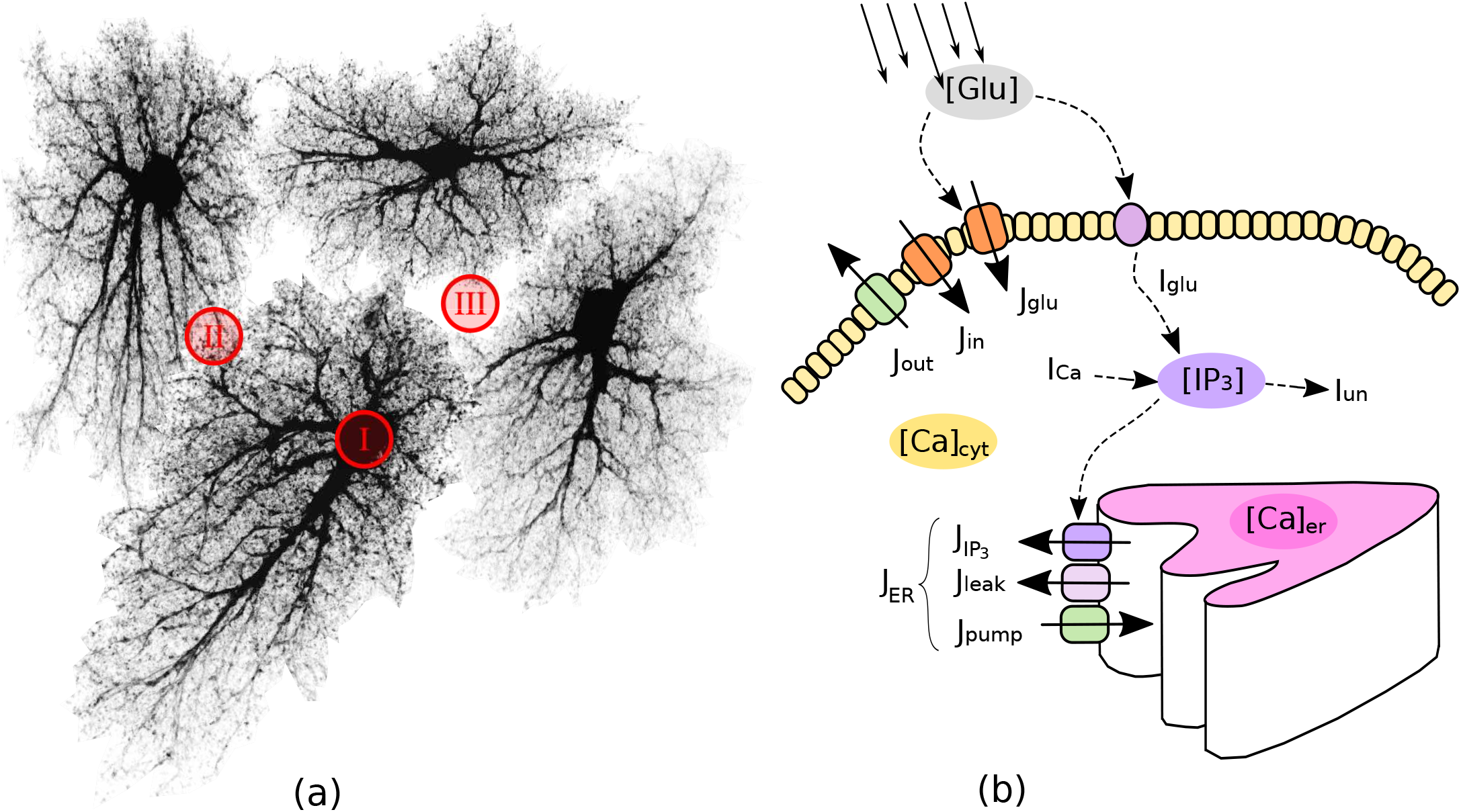
Model structure and molecular mechanisms. (a) Astrocytic network is segmented in three spatial compartments: I — cell bodies and thick branches; II—the mesh of thin branches; III — extracellular space. (b) Model variables (in grey ovals) and main regulatory pathways of intra-astrocyte calcium dynamics.

With regard to the local calcium dynamics, the extreme complexity and sheer number of cellular pathways involved, makes the detailed and comprehensive modeling of every Ca^2+^-related mechanism extremely challenging. Not surprisingly, there is a substantial body of published models that aim to account for the essential features of calcium dynamics in astrocytes, which do not completely agree with each other (Manninen and Havela, 2017). To choose the best model we build upon a model proposed by Ullah et al (2006) as a prototype, while other models could fit in the proposed approach as well, for example the “ChI” model by De Pitta et al (2009), which is similar to that of Ullah et al.

#### 2.1.1 Spatial structure

To represent astrocyte networks with realistic geometry of the regions I–III, one needs to create such templates algorithmically or, alternatively, obtain them from experimental data. Each of the two variants has its benefits and drawbacks. To provide just two examples, the experiment-based approach was employed in (Wallach et al, 2014) and an algorithmic creation of network templates was employed in (Postnov et al, 2009). Here we draw advantages from both approaches by suggesting a simple stochastic data-driven algorithm to create realistic surrogate spatial templates of astrocyte networks. Specifically, we use experimental images of astrocytes obtained from a public database, and arrange network structure using Voronoi partition and simple geometrical transformations.

#### 2.1.2 Neuronal activity

We assume that astrocytic Ca^2+^ response to local neuronal activity is primarily driven by the transporter-mediated uptake of neurotransmitters released from presynaptic membranes. One of the possible coupling mechanisms is the reversal of the Na^+^/Ca^2+^-exchanger transport due to an increase in [Na^+^] allowing for a Ca^2+^ influx. Here we sacrifice biophysical details in favor of model simplicity and assume that astrocyte calcium dynamics is excited directly by glutamate released from the presynaptic terminals, causing transient fluxes of Ca^2+^ through the plasma membrane. A typical cortex astrocyte is associated with 300–400 individual dendrites and is in contact with about 10^4^–10^5^ synapses (Bushong et al, 2002; Halassa et al, 2007). Judging by these numbers and taking into account sparsity of neuronal signaling in the cortex it seems safe to treat each pixel in the distributed model template as associated with a single or just a few individual synapses. For as long as we are not focused on information processing in the cortex, we can assume independent stochastic nature of spiking activity in any of the presynaptic units and describe local activity only statistically, neglecting any complex spike timing patterns. Consequently, we describe the synaptic glutamate drive to the model in each pixel as triggered by presynaptic spike trains drawn from independent homogeneous Poisson process *ξ*_*p*_(*t*) with intensity *p* Hz.

### 2.2 Astrocyte network topology

#### 2.2.1 Data-driven network generation

Astrocytes, like neurons, have complex morphology. Ideally, an algorithm to create spatial templates should provide means to “grow” realistic branching 3D shapes of astrocytes from a set of randomly placed “seed” locations. Indeed, there are many experimental and modeling studies of the branching patterns for various types of neurons (Cuntz et al, 2010; Donohue and Ascoli, 2008; Polavaram et al, 2014; Ascoli et al, 2007), providing means for creation of realistic surrogate shapes of as many neurons as needed. However, unlike neurons, there is less data available on the statistics of astrocyte branching, which makes it harder to create surrogate spatial templates of astrocyte networks. This hindrance can be circumvented by using a public database of microscopic images of cortical and hippocampal astrocytes (Martone et al, 2002, 2008).

To create a library of realistic spatial templates for individual cells, we downloaded a set of 27 fluorescent confocal 3D stacks of hippocampal astrocytes (4-week old rats, microinjection loaded with lucifer yellow in acute slices) (Bushong et al, 2004). Because our model is set in two-dimensional space, the stacks were flattened along the *z*-axis by max-projection, Figure 2. Each of the experimental astrocyte images then serves as a progenitor of randomized offsprings obtained by applying 250 random rotations (from 0^°^ to 360^°^, shearings and stretchings (within ± 20% of original size, uniform distribution), which results in a collection of 6750 randomized pseudo-experimental astrocyte templates, used to tile the model space. Such data set expansion from a limited number of “real-world” objects is a popular approach in machine learning (Simard et al, 2003; Krizhevsky et al, 2012) helping to prevent overfitting and providing for transformation-invariant feature learning.

**Figure 2.**
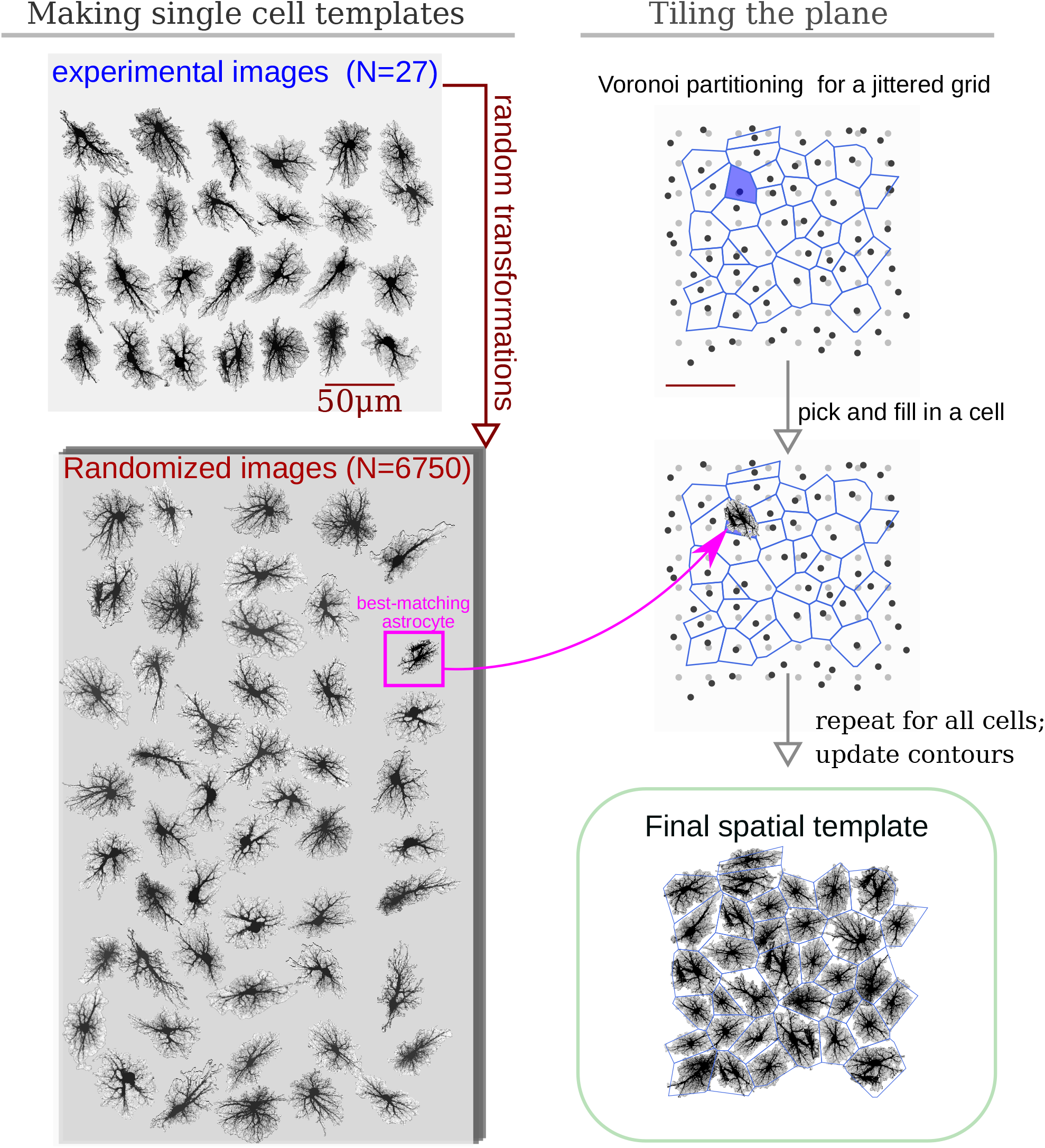
Algorithm to create surrogate templates of astrocyte network. First, a set of seeding points on a regular grid (light-gray) is perturbed with random shifts (dark-gray points). Voronoi diagram is then drawn for these points (blue lines). Each patch in the Voronoi partitioning is then filled with the best shape-matching template from an augmented collection of astrocyte images. The lookup collection is created from a set of experimental images taken from CCDB (Martone et al, 2008, 2002) by applying multiple different random rotations and shears to each experimental image.

Inspired by the fact that astrocytes establish distinct non-overlapping territories, we employ an algorithm based on Voronoi partitioning and active contours to tile the model space with astrocytes. First, we create a lattice of “seed points” regularly spaced at some intervals corresponding to average cell density, typically around 50 *µ*m, shown in light gray in Figure 2. The resulting regular grid is then deformed by jittering *x* and *y* coordinates of every point by a random displacement value drawn from Gaussian distribution with *σ* = 10 *µ*m (dark grey points in 2). Different values of spacing and jitter can be used, the ones used here tended to give the most realistic tiling results. Next, a Voronoi diagram, which for each seed point delineates territories closer to it than to any other seed point, is drawn for the jittered points. We then iteratively pick a polygonal area patch from the Voronoi partitioning, look up a template astrocyte from the randomized collection, with a convex hull best matching the shape of the given Voronoi patch, and place this template into the model space. Repeated for all patches in the Voronoi partition, this creates a preliminary tiling with partially overlapping domains of neighboring astrocytes and occasional empty spaces. Next, this draft tiling is optimized with an active deformable model: the perimeter of each cell template is treated as an elastic two-dimensional curve, which optimizes an energy functional designed to promote repulsion between overlapping regions and adhesion between neighboring cells, with a penalisation of the major cell shape deformation. After all domain boundaries are settled, the spatial templates are interpolated into the deformed contours. The described process of the network template creation is visualized in Supplementary video 1.

#### 2.2.2 Computational design

Our simulations are based on compiling an encoded raster image representation (a template) of the model space to region-specific equations. For the sake of computational simplicity, we use two-dimensional spatial layout — each pixel of the spatial template can be interpreted as a thin slab, occupied either exclusively (e.g. in the soma) or partly by astrocyte cytosol; or as belonging to extracellular space. As follows from this approach, each pixel in the model space has to be assigned to either astrocyte-free space (class III) or astrocyte-occupied space, ranging from class I, astrocyte soma and thick branches, to class II, i.e. elements of volume containing a tangle of thin astrocyte processes and unresolved neuronal structures, e.g. synaptic boutons. We account for a graded transition from thick branches to thin processes to leaflets by introducing a local astrocyte volume fraction (AVF) parameter, which defines the landscape of how much of each pixel volume is occupied by astrocyte in the 3D prototype (Figure 3 A–B). To describe the relative input of the store-operated calcium flux and plasma membrane flux, we introduce a surface-volume ratio (SVR) parameter, which inversely depends on AVF (Figure 3 C). The SVR value is maximal at the edges of the leaflets and minimal in the soma. Accordingly, a simple raster image serves as a spatial template to encode the model space. Specifically, non-zero values in the blue channel define astrocyte-occupied pixels, while intensity in the red channel encodes AVF and ranges from minimal value (class II) to 1 (class I). Thus, one can set up computation for a specific spatial template by simply drawing it algorithmically or with an indexed palette using a graphical editor.

**Figure 3.**
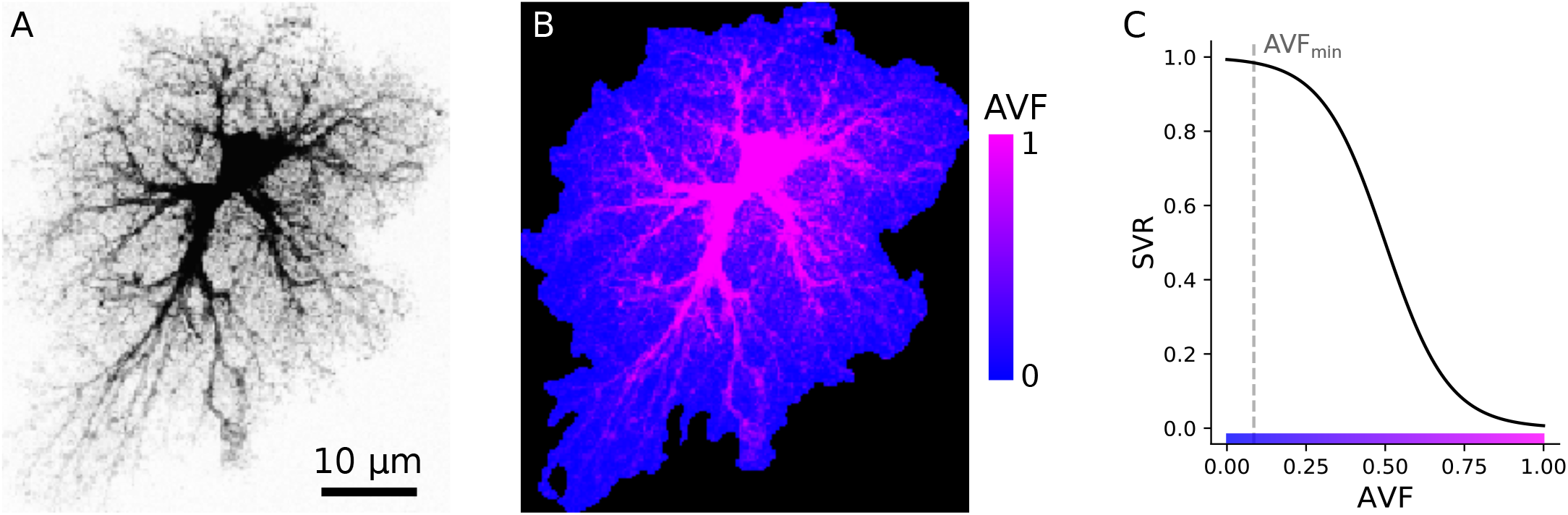
Example of AVF color-coding. (A) Maximal projection of a confocal image of a cortical astrocyte. (B) color-coded template ready for simulation; regions with non-zero blue channel delineate astrocyte domain, while intensity of the red channel encodes AVF. (C) Link between color-coded AVF and SVR parameters.

At each integration step the master program module optionally compiles the provided image into a set of equations by mapping each pixel color to equation set following the color-coded dictionary. For each pixel first the point dynamics is applied, i.e. right-hand terms are evaluated. Next, possible substance exchange and short-range interactions are taken into account based on the class of the neighboring pixels. This approach is flexible, but has an overhead of compiling the color-to-equation mapping. To improve the computational performance we employ NVIDIA CUDA, a parallel GPU-based computing technology. Numerical integration of the model differential equations is done in an explicit scheme (4th order Runge–Kutta method adopted for stochastic differential equations) implemented in AGEOM–CUDA software (Postnov et al, 2012).

### 2.3 Intracellular calcium dynamics: principal quantities and flows

The model for local Ca^2+^ dynamics is based on that of Ullah et al (2006) with a few modifications previously introduced in (Brazhe et al, 2018), which sum up to treating ER calcium as a dynamic variable, adding neurotransmitter-dependent calcium influx via plasma membrane, and segregation between thick and thin processes. Below we describe the proposed model, focusing on the differences with the Ullah et al (2006) model; equations and parameters that are the same here as in the Ullah model are omitted.

The principal variables of the model are (i) the cytosolic calcium concentration [Ca^2+^]_*c*_, (ii) calcium concentration in the endoplasmic reticulum [Ca^2+^]_*ER*_, (iii) inositol trisphosphate concentration in the cytosol [IP_3_] and (iv) extracellular glutamate concentration [Glu].

To account for the morphology-based spatial heterogeneity of astrocytes, we introduce a parameter *r* ∈ (0 *< r*_min_ … 1] — a scalar quantity, roughly representing local AVF. This parameter also defines a linked parameter *s* representing local SVR of the astrocyte processes; SVR is inversely related to AVF: 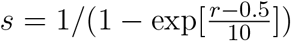. SVR scales relative inputs of Ca^2+^ exchange through plasma membrane and with ER as described below, while AVF scales effective diffusion coefficients for Ca^2+^ and IP_3_.

The equation set for the principal variables reads:

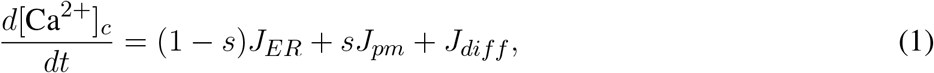

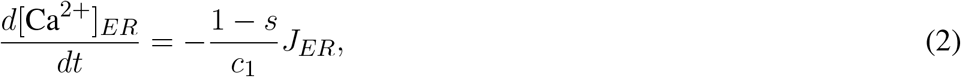

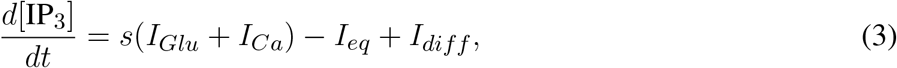

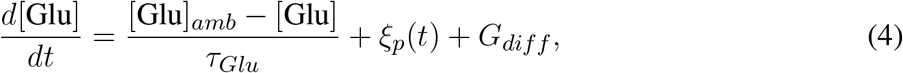

where *J*_*ER*_ is the total flow of calcium ions in exchange between the cytosol and endoplasmic reticulum; *J*_*pm*_ is the total flow of calcium ions through the plasma membrane in exchange between the cytosol and extracellular space; *I*_*Glu*_ and *I*_*Ca*_ stand for glutamate- and calcium-dependent inositol trisphosphate production mechanisms, mediated by phospholipases *β* and *δ*; *I*_*eq*_ is a simplified first-order equilibration of inositol trisphosphate concentration to the basal level [*IP*_3_]_0_; [Glu]_*amb*_ is the ambient concentration of extracellular glutamate, and *τ*_*Glu*_ is the timescale of its clearance and return to the baseline level; *ξ*_*p*_(*t*) isstochastic source of glutamate from nearby located neuronal synapses triggered by Poisson spike trains in each pixel. Additionally, *J*_*diff*_, *I*_*diff*_, and *G*_*diff*_ describe the finite-element approximation of diffusion of cytosolic Ca^2+^, IP_3_ and extracellular glutamate, respectively and are described below.

The weighting coefficient *s* accounts for the stratification of intracellular dynamics according to Figure 3: in the leaflets *r* = *r*_min_ ≪ 1 and *s* ≈ 1, while for deep cytosol locations *r* = 1 and *s* ≈ 0. With this we (i) describe that input from all plasma membrane calcium currents is maximal in the leaflets and (ii) assume that endoplasmic reticulum doesn’t invade leaflets much, thus ER exchange is large only in thicker branches and soma. We also assume that all IP_3_ is produced in the plasma membrane by means of G-protein coupled phospholipase *β* or Ca^2+^-dependent phospholipase *δ*.

#### 2.3.1 Calcium exchange between the cytosol and ER

Total calcium flow across the endoplasmic membrane is composed of IP_3_R-mediated current 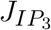, leak of Ca^2+^ from endoplasmic reticulum *J*_leak_, and contribution of ER membrane Ca^2+^ pump *J*_pump_:

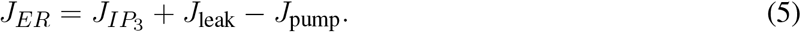

Ca^2+^ current via IP_3_ receptors 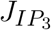 is modelled in the same way as in Ullah et al (2006) (equations 2, 4, 5, 9–12). Two other terms in (5) stand for the leak of calcium from *ER*, and for the pumping it back, respectively, following equations (3,6) in Ullah et al (2006).

#### 2.3.2 Transmembrane calcium flows

Transmembrane calcium exchange *J*_pm_ consists of three flows:

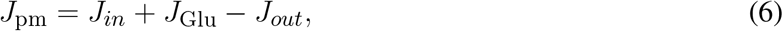

where *J*_*in*_ describes the sum of background constant Ca^2+^ influx and agonist-dependent IP_3_-stimulated Ca^2+^ influx across plasma membrane from the extracellular space and *J*_*out*_ is an extrusion current (eqs. 7–8 in Ullah et al); *J*_Glu_ = *γ*[Glu] describes the direct effect of extracellular glutamate on additional calcium influx.

#### 2.3.3 Inositol trisphosphate turnover

The dynamics for IP_3_ concentration (3) has the following terms: first, we use a lumped first-order description of [IP_3_] equilibration to a resting level [IP_3_]_0_:

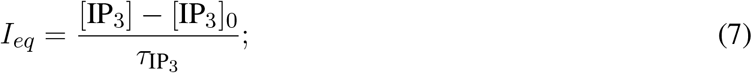

second, we account for the Ca^2+^-stimulated IP_3_ production in the same way as in Ullah et al (2006), (eq. 14), and third, we account for glutamate-driven IP_3_ production *I*_*Glu*_ following Ullah et al (2006), (eq. 15).

#### 2.3.4 Synaptic glutamate drive

Stochastic glutamate source *ξ*_*p*_(*t*) in each pixel is modeled as quantal release triggered by a spike train drawn from a homogeneous Poisson process with intensity *p*, which agrees with statistics of neuronal firing (Softky and Koch, 1993). Accordingly, the *ξ*_*p*_(*t*) term in (4) is given by

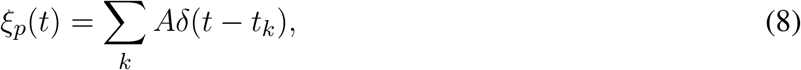

where *A* is the instantaneous increase in glutamate release rate associated with each presynaptic event and *t*_*k*_ are times of presynaptic spikes in the given pixel following Poisson process with intensity *p*_*syn*_.

### 2.4 Intracellular diffusion

Elevated cytoplasmic Ca^2+^ can remain confined to the spatial domain of a single astrocyte, but can also spread to the neighboring astrocytes (Falcke, 2004; Carmignoto, 2000; Nedergaard, 1994) in a wavelike manner. The involvement of a large number of cells into a wave is still not fully understood though it may be an important aspect of information processing in the brain (Haas et al, 2006). In addition to the described above glutamate-mediated neuroglial interaction, other spatial coupling pathways, such as the release of extracellular ATP (with its degradation to ADP) or release of another neurotransmitter, GABA, (Serrano et al, 2006; Newman, 2003; Bowser and Khakh, 2004) are known but have not yet been studied in detail from the viewpoint of the resulting network behavior. Thus, at least two main mechanisms can account for the intercellular wave propagation: (i) diffusion of *extracellular* ATP and its action on P2Y receptors on astrocytic membranes and (ii) diffusion of *intracellular* IP_3_ and Ca^2+^ between contacting astrocytic leaflets, via gap junctions. Relative input of the two mechanisms differs across brain regions and for the cortical astrocytes the one mediated by the gap junctions has been reported to prevail (Haas et al, 2006). The current work is therefore focused on the latter mechanism. Accordingly, astrocytes in the model are networked by an analog of gap junctions dispersed over the parts of the cell perimeter and simulated as the connection of areas with low AVF.

Here we employ a rather simplified description of diffusion in the cytoplasm where the region occupied by astrocyte is considered as a continuous space. As corroborated by evidence for autologous gap junctions between the processes of the same astrocyte (Wolff et al, 1998; Nagy and Rash, 2003; Genoud et al, 2015), astrocyte cytosolic volume can be described as a porous sponge-like medium rather than a branched structure or acyclic graph. Thus, possible hindrance to IP_3_ or Ca^2+^ diffusion through the intricate mesh of astrocytic processes due to tortuosity and porosity of the astrocytic volume can be accounted for by a simple scaling of the apparent diffusion coefficient. Though an interesting issue, a detailed account for intracellular diffusion and connectivity between neighboring points in an astrocyte is outside the scope of the current study and here we resort to a rather minimalistic description.

We account for Ca^2+^ and IP_3_ diffusion throughout astrocyte as exchange between neighboring astrocyte-containing pixels. This also includes exchange at borders between neighboring astrocytes to imitate the function of gap junctions, which is supported by evidence that IP_3_ can diffuse through the gap junctions along with Ca^2+^ (Yule et al, 1996).

Specifically, the diffusive term in (1), e.g. for Ca^2+^ reads:

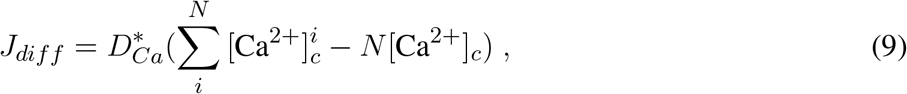

where 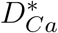 is the diffusion rate defined as the diffusion coefficient for Ca^2+^ scaled by spatial grid step *δx* = 0.59 *µ*m/pixel and local AVF value *r*: 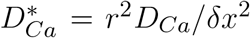, and *i* enumerates the *N* nearest neighboring astrocyte-containing units. Here we regard diffusion in porous media and assume that larger AVF is associated with larger cross-sectional area open for diffusion, as the astrocyte process diameter increases, and simultaneously with less tortuous paths taken up by diffusing molecules as the processes become less entangled. This leads to approximately quadratic scaling of *D*^∗^ with *r*. The diffusive term for IP_3_ is defined in a similar way.

Finally, neighboring pixels with different AVF values obviously contain unequal volumes of astrocyte cytoplasm; hence, small concentration changes in areas with high AVF should cause larger diffusion-mediated concentration changes in the neighboring pixels with low AVF. This was accounted for by scaling the concentration rates of change due to diffusive exchange by the ratio of the AVF values of the two neighboring pixels.

### 2.5 Model parameters

The basic set of model parameters is given in Table 1. Only new parameters and values different from that in Ullah et al (2006) are shown. The few values that are different were adjusted in order to provide the reasonable dynamics with the introduced treatment of [Ca^2+^]_*ER*_ as a variable in our model.

**Table 1.** Model parameters

| Param. | Value | units | Remarks |
| --- | --- | --- | --- |
| $c_0$ | not used | | Total $[Ca^{2+}]$ in terms of cytosolic vol |
| $v_4$ | 0.1 | $\mu M/s$, | max rate of $Ca^{2+}$-stimulated $IP_3$ production |
| $v_5$ | 0.01 | $\mu M/s$, | rate of $Ca^{2+}$ leak across plasma membrane |
| $k_1$ | 1.0 | 1/s, | rate constant of $Ca^{2+}$ extrusion |
| $k_3$ | 0.05 | $\mu M$, | activation constant for $Ca^{2+}$-pump |
| $\tau_{Glu}$ | 0.1 | $\mu M/s$ | rate constant for synaptic glutamate uptake |
| $p_{syn}$ | 0.005–0.01 | Hz | rate of Poisson process, for glutamate release |
| $A$ | 27 | $\mu M$ | amount of quantal glutamate release |
| $[Glu]_{amb}$ | 0 | $\mu M$ | ambient extracellular glutamate |
| $D_{Ca}$ | 10 | $\mu m^2/s$ | diffusion coefficient for cytoplasmic $Ca^{2+}$ |
| $D_{IP_3}$ | 10 | $\mu m^2/s$ | diffusion coefficient for cytoplasmic $IP_3$ |
| $D_{Glu}$ | 2 | $\mu m^2/s$ | diffusion coefficient for extracellular glutamate |
| $r_{min}$ | 0.085 | dimensionless | minimal value of the AVF |

## 3 RESULTS

The proposed model, including the modifications to the local calcium dynamics and spatial mapping, was tested in a number of simulation experiments with different parameter settings and different spatial templates (which we call “cells” for shorthand below). To test for agreement between model behavior and the experimentally observed dynamics, first, we looked at the effect of the level of mean neuronal firing rate (local rate of the Poisson point process in terms of our model) on spatio-temporal dynamics of astrocytic calcium, and second, we tested whether the artificial spatial templates could provide for realistic intercellular calcium waves or other collective variants of astrocytic calcium dynamics.

### 3.1 Wave patterns in single-cell templates

Figure 4 summarizes simulations of the 27 single-cell templates shown in Figure 2. At low excitation (*p*_*syn*_ = 0.005 Hz) most Ca^2+^ events were spatially confined and tended to start at a small number of sites, as shown with max-span contours and red labeling in Figure 4 (panel A, left) for 25 largest events. At higher excitation (*p*_*syn*_ = 0.01 Hz) many Ca^2+^ events spread to occupy the whole cell domain and again tended to initiate at the same sites. Synaptic signaling events were integrated into a spatial glutamate profile as shown in Figure 4 (panel A, bottom): local surges of extracellular glutamate are sparse at low excitation, while their instantaneous spatial density increases at high excitation, with a tendency of nearby sparks to blend.

**Figure 4.**
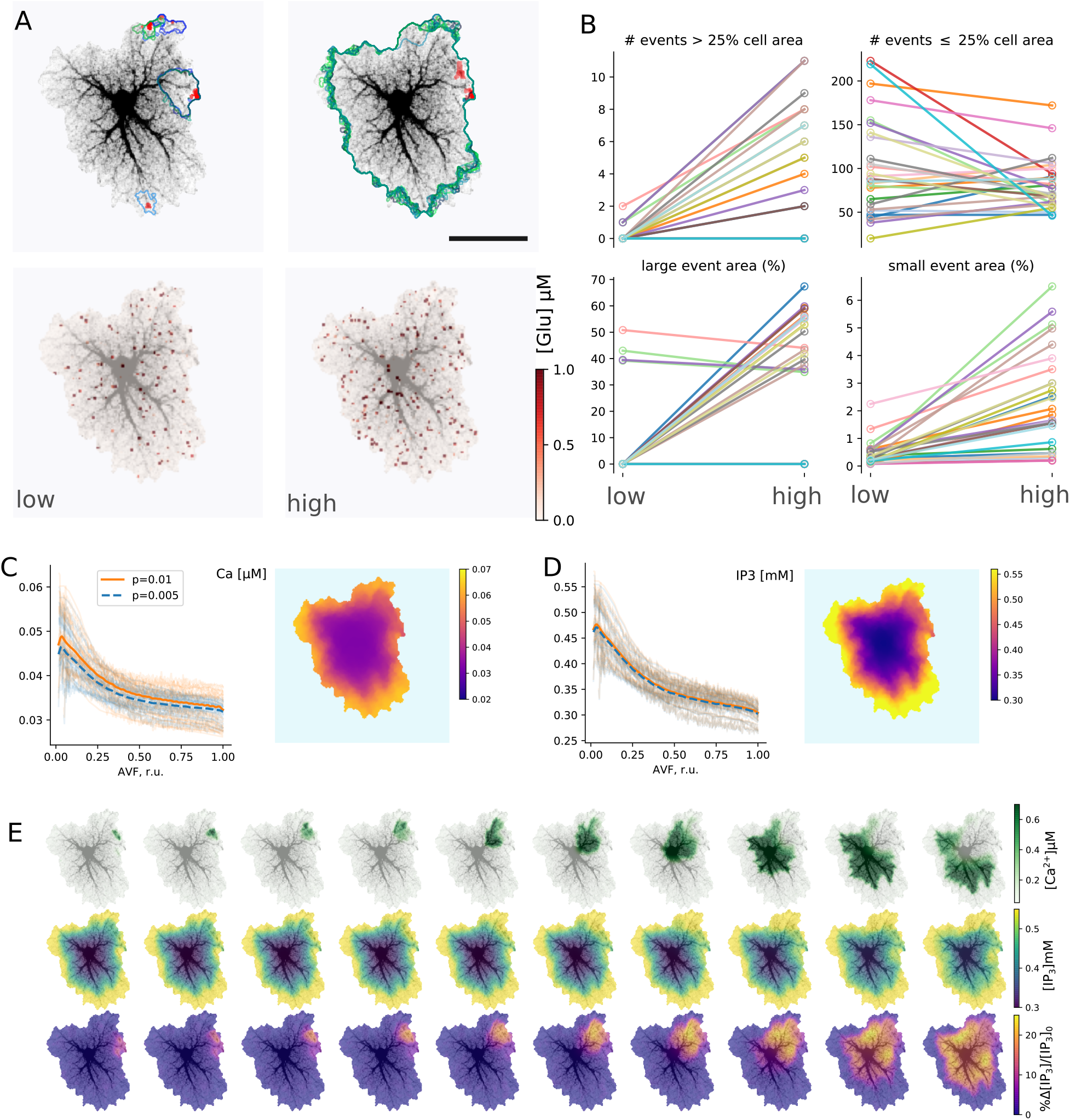
Simulations of single-astrocyte spatial templates. (A) top row: Ca^2+^ transient initiation sites (red) and maximum span contours at low (left, *p*_*syn*_ = 0.005 Hz) and high (right, *p*_*syn*_ = 0.01 Hz) synaptic drive, in both cases 25 largest events are shown, at high drive all such events span the whole template; bottom row: snapshots of instantaneous extracellular glutamate concentrations at low and high synaptic drive parameters. Scale bar: 25 *µ*m. (B) Effect of synaptic drive on Ca^2+^ transient frequency and sizes (*n* = 27 templates, simulation time 2500 s after burn-in period of 2000 s); top row: number of events covering more than 25% of cell area (“large” events) increases with excitation strength (left), number of events covering less than 25% of cell area (“small” events) decreases (right); bottom row: average areas of both “large” (left) and “small” (right) events increases with excitation strength in most templates. (C) Distribution of baseline [Ca^2+^]_*i*_ levels with AVF parameter at low and high stimulation drives (left) and an example of spatial distribution (right). Each transparent line corresponds to one template, thick lines: average. (D) Same as in (C), but for [IP_3_]_*i*_. (E) Evolution of a single Ca^2+^ transient starting in top-right corner of the template; top row: [Ca^2+^]_*i*_, middle row: [IP_3_]_*i*_, bottom row: relative change in [IP_3_]_*i*_.

The tendency, exemplified by a single template in Figure 4 A, was supported by the majority of single-cell templates (Figure 4 B): the number of events (during 2500 s simulation time) covering more than 25% of the cell area increased with excitation for nearly all cells except three, which were incapable of generating whole-cell transients. Most cells demonstrated a decline in the number of small events, covering less than 25% of the cell area with excitation, as a larger proportion of events was enabled to spread over larger areas, while the rate of event initiation could remain stable. The area of large (*>* 25%) events increased for all cells which generate large events under low drive conditions except the three, which didn’t generate large events at all, apart from other cells. The average area of the small (*<* 25%) events increased with excitation strength for all cells.

Stochastic local glutamate surges initiate two parallel processes: fast localized Ca^2+^ transients and slower IP_3_ production. Both integrate over time to steady-state levels of the model variables. Because the relative input of plasma membrane transport is defined by AVF in our model, we expected that the steady state levels and the probability of Ca^2+^ event initiation should depend on AVF as well. Steady-state values of [Ca^2+^] and [IP_3_] decreased with growing AVF (Figure 4 C–D), forming an uneven spatial profile. Calcium, as well as IP_3_ levels, were higher in the periphery and lower near the soma, which is due to Ca^2+^ entry during the synaptic excitation and due to higher IP_3_ production in the regions with lower AVF. Different cells varied in steady-state levels of the variables, while intensified stimulation lead on average to a slight elevation of steady-state [Ca^2+^]_*i*_, due to increased Ca^2+^ entry and didn’t affect [IP_3_]_*i*_, as the increase in [Ca^2+^]_*i*_ was insufficient for activation of PLC_*δ*_.

An example of a single large Ca^2+^-event is shown in Figure 4 E. The expanding wave of elevated [Ca^2+^]_*i*_ initiates in a small location and spreads over the whole spatial domain (top row). Due to the large range of steady-state [IP_3_]_*i*_ concentrations, the event is unclear in absolute [IP_3_]_*i*_ values (middle row), but is obvious in the relative scale (bottom row).

Despite the stochastic nature of excitation, Ca^2+^ activity in most cells self-organized in a repeated pattern of Ca^2+^ transient initiation and spreading (Figure 5); event initiation sites were tightly clustered. Interestingly, activation in some clusters lead to spatially confined events, unlinked to activity in the rest of the cell, while transients originating in other sites tended to spread over the whole spatial domain in a repeated fashion. This is illustrated in Figure 4 (A–D) for an example spatial template (see also Supplementary Video2). This cell is markedly anisotropic, thus shaping the dominant wave spreading properties. The two active initiation sites labeled as #1 and #2 display different properties: events, starting in the site #1 often spread over large portions of the cell, as shown for a line-scan path in (B), while line-scan along the path starting in #2 was either activated by a wave coming from #1 or — very locally and with a higher frequency — by confined transients initiating in #2. Averaging small temporal windows around Ca^2+^ spikes at the origin of path #1 shows that activation along this path is time-locked to activation of the initiation site. On the other hand, averaging similar temporal windows triggered by Ca^2+^ spike at the origin of path #2 didn’t reveal any structured activation patterns. Figure 5 (D) shows max-span contours of 25 largest events with their initiation sites mapped in red, as well as 5 peak-delay maps for a repeated pattern of activation. In these maps color indicates delay in seconds between the Ca^2+^ peak at the initiation site and the Ca^2+^ peak at each point of the cell. The precise wave initiation site could vary within 3–5*µ*m, but the overall spatiotemporal pattern remained similar, with activation spreading mainly along the long thick processes towards the soma. Additional examples of repeated patterns for other cells are provided in panel (E). In some cells preferred wave initiation sites could alternate between two polar positions or a few neighboring regions. There was also some scatter in the maximum delay between the initiation of the wave and its full expansion.

**Figure 5.**
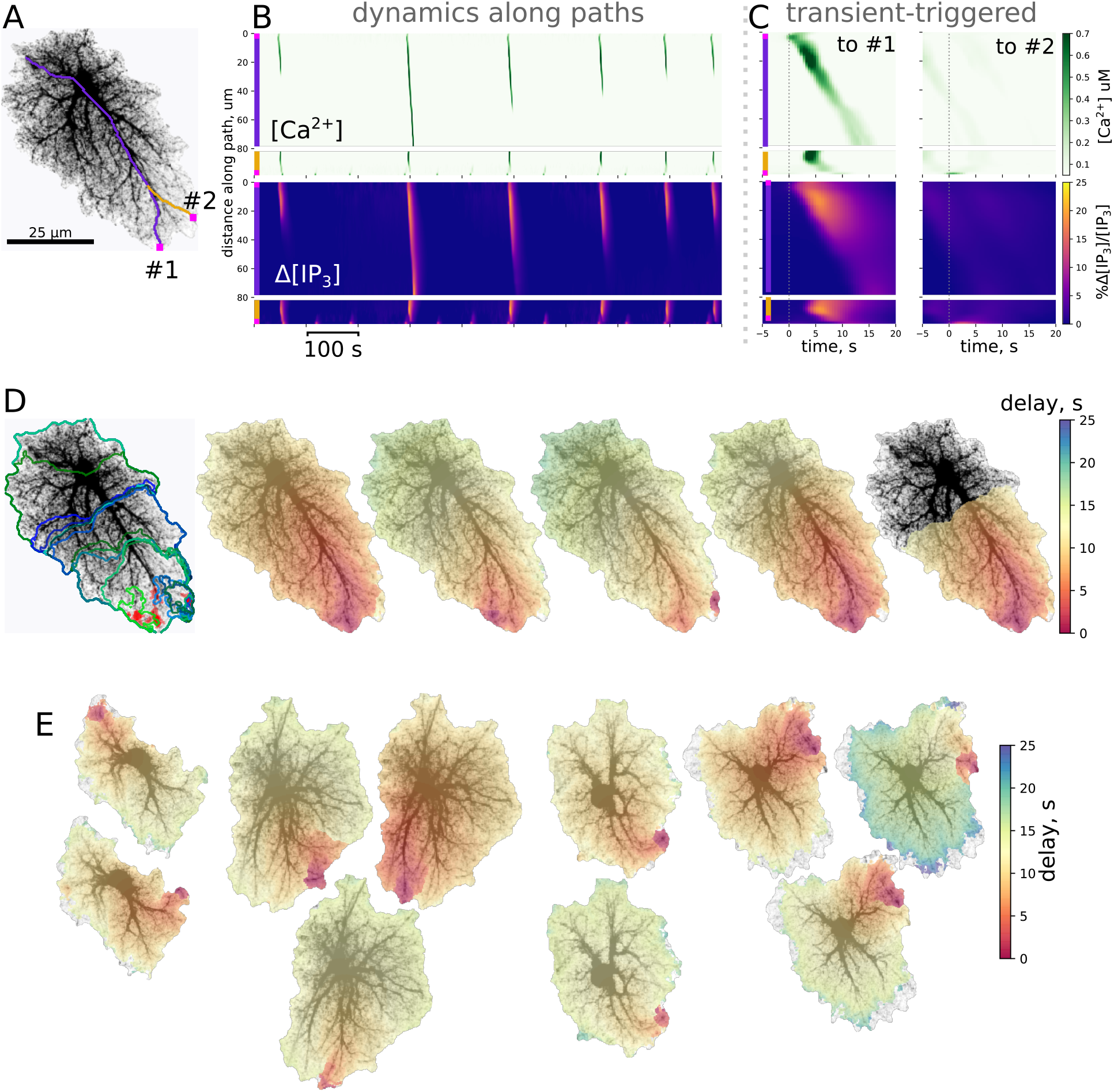
Ca^2+^ dynamics in single-cell templates self-organizes in repeatable spatiotemporal patterns. A spatial template for simulation and 2 path-scans (#1, purple and #2, yellow) used for rasters in; (B) Rasters of [Ca^2+^]_*i*_ and relative change in [IP_3_]_*i*_ along paths; path starting points are shown as magenta squares. (C) Transient-triggered averages, linked to the [Ca^2+^]_*i*_ peaks at the origin of path #1 (left) and to the [Ca^2+^]_*i*_ peaks at the origin of path #2 (right); initiation of a [Ca^2+^]_*i*_ peak at the origin of path #1 typically leads to a full-cell [Ca^2+^]_*i*_ wave, while [Ca^2+^]_*i*_ transients emerging at the origin of path #2 remained localized. (D) Left: clustered initiation points (red) and contours of 25 biggest [Ca^2+^]_*i*_ waves; right: 5 “large” [Ca^2+^]_*i*_ waves with color-coded delay before reaching the peak in [Ca^2+^]_*i*_ show a tendency to start in the same area and spread with similar spatiotemporal profiles. (E) Examples of repeated spatiotemporal patterns in 4 other spatial templates: in some cells there was more than one preferred initiation site, the full delay before initiation and waning of the wave also varied.

Though localised Ca^2+^ events could be initiated in the low-AVF regions due to direct influx through the plasma membrane, these event seeds needed to reach a tip of a thicker branch with higher AVF to be amplified by IP_3_-mediated ER exchange. Thus, initialization of a global Ca^2+^ wave critically depends on a coincidence of exactly the right spot in the AVF profile — allowing both for a high enough [IP_3_]_*i*_ baseline and sufficient ER exchange — and a wide and long enough cluster of glutamate release due to local increase in synaptic activity. Areas containing only thin processes with low AVF will display only frequent local Ca^2+^ sparks, unable to invade the neighboring regions and thus will primarily set the baseline levels of [Ca^2+^]_*i*_ and [IP_3_] due to diffusion. Indeed, astrocytes in hippocampal slices display frequent localized Ca^2+^ events in the cell periphery, often termed “microdomains”, with a characteristic size of high-Ca^2+^ spots much smaller than the cell size and originating in the thin processes region (Rungta et al, 2016). At the same time, the “lifespan” of the Ca^2+^ wavefront increases with the already invaded area of the thick branch region due to regenerative Ca^2+^ release, which in turn relies on background [IP_3_] level. Thus, the specific topology of the cell template predicts the “hot spots” for the probability of Ca^2+^ wave seeding.

To quantify spatiotemporal properties of the simulated Ca^2+^ dynamics, we examined complementary cumulative distribution functions (CCDF) of areas and lifetimes of individual Ca^2+^ transients in all spatial templates to avoid selection bias. To this end, we thresholded and segmented the simulated calcium dynamics at 25% change from the local baseline level and treated the resulting contiguous *TXY* volumes of suprathreshold Ca^2+^ as discrete events. The left column of Figure 6 provides statistics of areas, covered by each event and shows the probability *P* that the area *S* occupied by a calcium elevation event is equal to or greater than *s*: *P* (*S >*= *s*). The right column describes durations of calcium events and shows the probability *P* that the lifetime *T* of the event exceeds *t*: *P* (*T >*= *t*). Red curves correspond to the case of low neuron activity, the blue ones correspond to high neuron activity. In both cases the CCDF curves are presented in double logarithmic coordinates and can be approximated by a straight line within some ranges of event areas or durations, which implies a power-law behavior and agrees with experimental data (Wu et al, 2014). If a given variable is distributed according to a power law with probability density function (PDF) *p*(*x*) ∝ *x*^−*α*^, then the CCDF also has a power-law behavior, but with a smaller exponent *P* (*x*) ∝ *x*^−(*α*−1)^. After recalculating from CCDF to PDF exponents, the resulting parameters were different from that reported in (Wu et al, 2014): for areas, *α* ≈ 3.3 … 4.0 in the model versus *α* ≈ 2.1 2.4 in cultured astrocytes, and for durations *α* ≈ 4.5 … 5.1 in the model versus *α* ≈ 1.97 … 2.16. These discrepancies can be explained by imperfection of the model and the 2D spatial embedding in the model. Defining model parameters that govern the shape of the event size and duration distributions is a potentially interesting outlook for further studies. Increase in synaptic excitation favored larger areas occupied by waves and longer durations of each event. The kinks in the CCDFs for event areas correspond to transition to whole-cell waves, i.e. events covering more than 30–50 *µ*m^2^ were likely to expand further and cover the whole cell.

**Figure 6.**
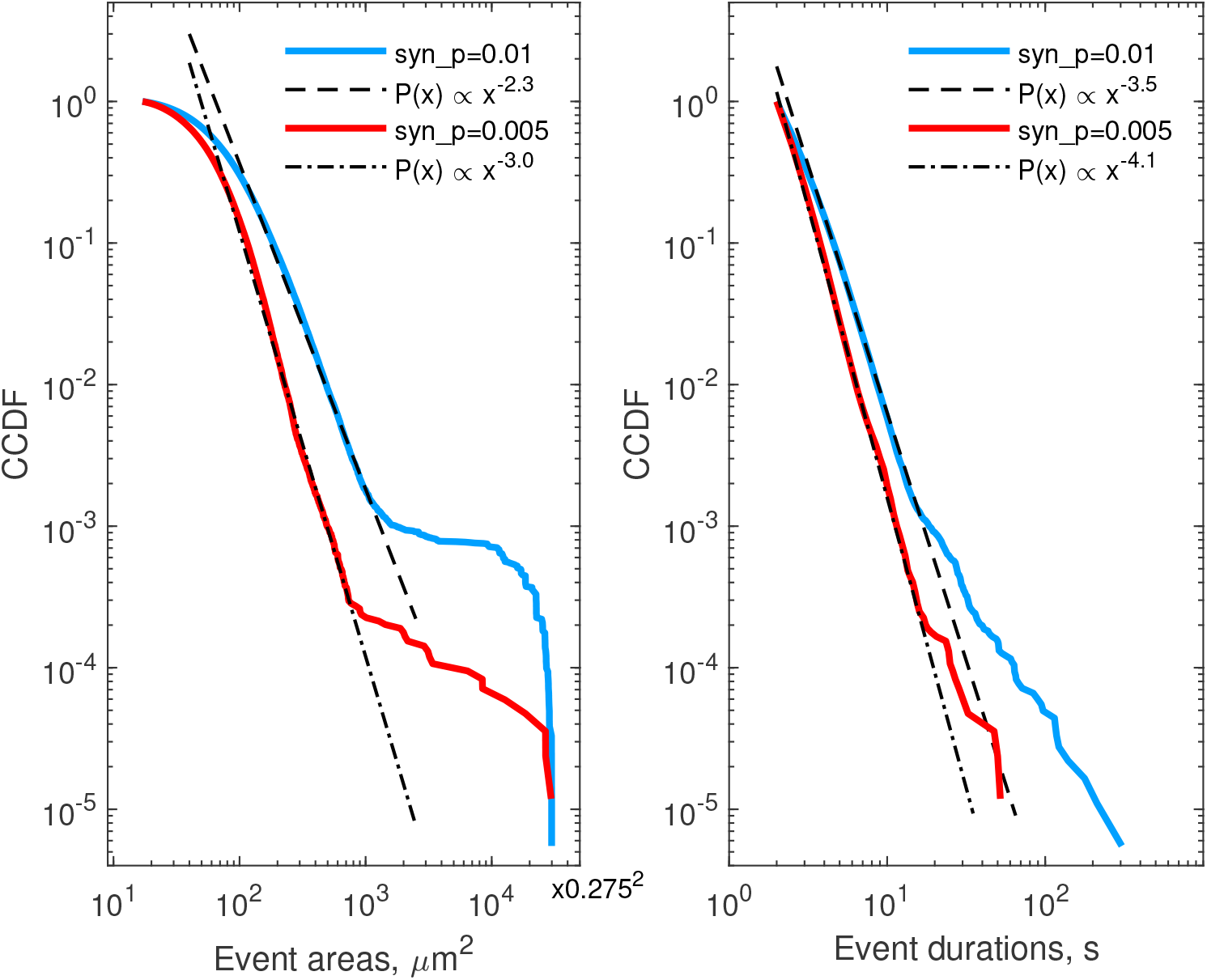
Complementary cumulative distribution functions for areas (left) and durations (right) of Ca^2+^ events in all single-cell templates. Red lines — low excitation (*p*_*syn*_ = 0.005 Hz), blue lines — high excitation (*p*_*syn*_ = 0.01 Hz). Event area CCDFs: slopes of the fits for the low and high excitations are 3.0 and 2.3, correspondingly; the bend at the large areas corresponds to transition to whole-cell activation. Event duration CCDFs: slopes of the fits are 4.1 and 3.5 for the low and high excitations, correspondingly.

### 3.2 Collective dynamics of astrocytes

After testing model behavior at microdomain and single-cell level, we turned to ensembles of connected cells. Collective dynamics of astrocytes was simulated using spatial templates containing about 40 cells as shown in Figure 7 (A); see also Supplementary Video 3. The simulated network is smaller than the size of cliques or networks of connected astrocytes in the neocortex (Houades et al, 2008), but still at the same order of magnitude. Simulations of larger spatial network templates are also possible but are more computationally demanding. Ca^2+^ activity was not uniform across the spatial template: there were active “hot” brims in the periphery of most of the cell domains and some cells were less active than others – Figure 7 (B). We observed Ca^2+^ transients originating in some astrocytes and expanding to their neighbors in a wavelike manner, an example of such a wave is shown in Figure 7 (C) as a sequence of contours of elevated [Ca^2+^]_*i*_, separated by 1 s.

**Figure 7.**
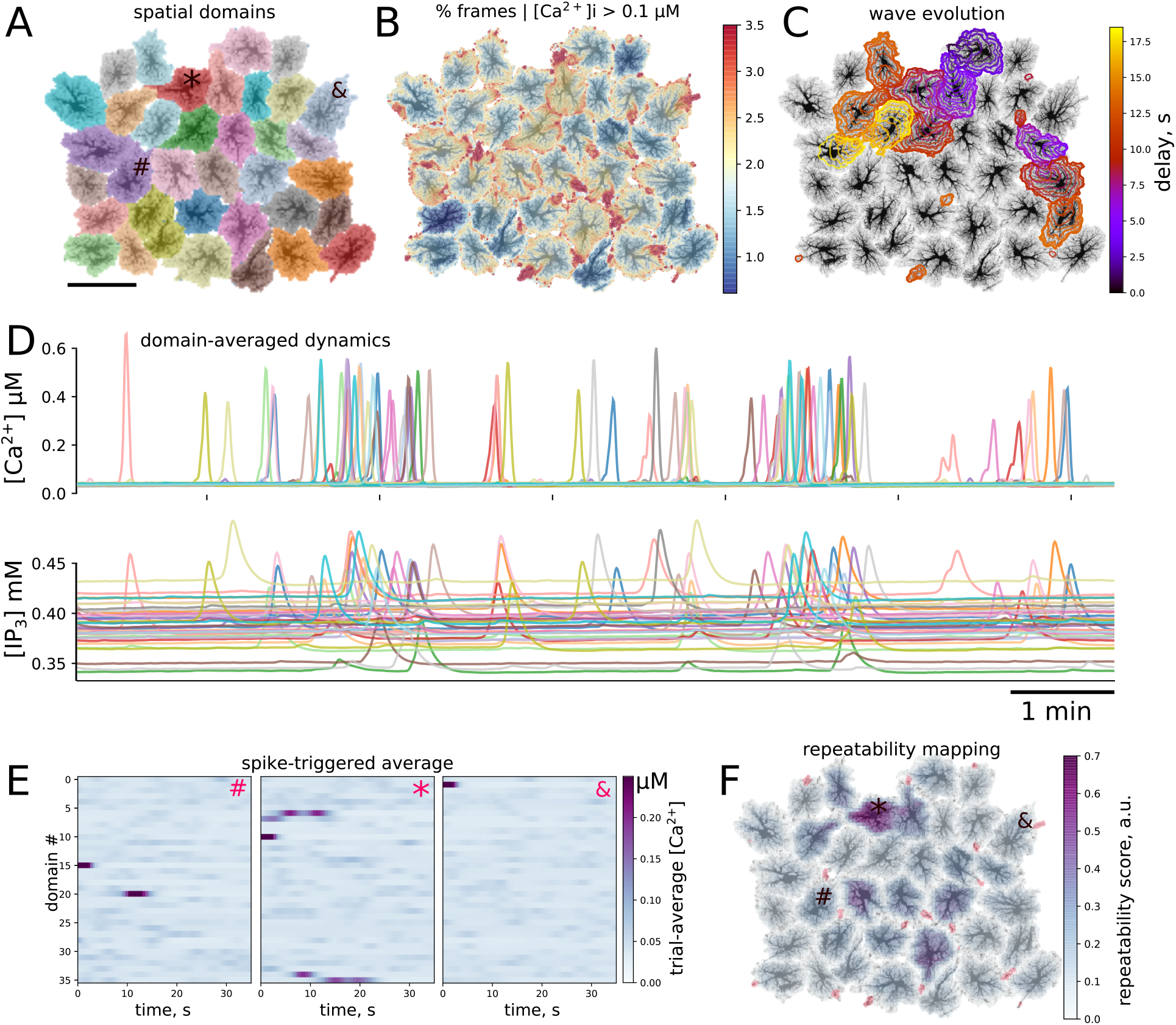
Ca^2+^ activity in multicellular template. (A) Layout of individual spatial domains, each cell is color-coded. (B) Activity level is not uniform: color-coded percentage of time that each pixel had [Ca^2+^]_*i*_ above 0.1*µ*M. Domain periphery is more active than somatic regions. (C) Example of spatiotemporal evolution of two co-occuring waves. Contours are separated by 1 s, time delay since the first contour is color-coded. (D) Cellwise dynamics: Ca^2+^ and IP_3_ values averaged over individual cell domains, line color corresponds to the map in (A), visible are single-cell events as well as packed Ca^2+^ transients representing multi-cellular waves. (E–F) Repeated patterns of cell activation. (E) Spike-triggered averages of cellwise [Ca^2+^]_*i*_ profiles initiated by the cells indicated as #, *, and & in (A) and (F); activity of the first cell is time-locked to activation of a single other cell, Ca^2+^ transient the second cell consistently leads to activation of several cells, while the third cell doesn’t participate in repeatable patterns. (F) Local score of activation pattern repeatability based on approach shown in (E) (see text for more details); activation of a subpopulation of cells leads to repeated activation of its neighbors; red regions denote wave initiation sites as in Figure 4 (A).

To simplify the description of emerging spatiotemporal patterns, each cell can be considered as an element that “fires”, producing a global whole-cell spike, or remains silent. This cell-wise activity is shown for [Ca^2+^]_*i*_ and [IP_3_]_*i*_ in Figure 7 (D), where line colors correspond to domain colors in panel (A). We observed individual cell spikes as well as packs of tightly grouped spikes corresponding to multi-cellular waves. A considerable scatter is clear in the baseline levels of IP_3_, which reflects individual properties of each cell. IP_3_ concentration peaks are wider and occur with a small delay in comparison to Ca^2+^ peaks, which reflects the slower kinetics of IP_3_ production timescale and a lag due to diffusion of IP_3_ from the periphery to the somatic region.

A large Ca^2+^ transient in one astrocyte can spread to other cells. We were interested if the networked spatial templates would also develop repeated patterns of activation. As a simple test for repeatable patterns, we calculated averages of the domain-averaged Ca^2+^ rasters in a short time window, triggered by Ca^2+^ spikes in different domains. In the case of stable activation sequence, the spikes appear with the same time-lag relative to the seeding astrocyte, and are thus visible in the raster plot, while if spiking in other astrocytes is not time-locked to the seeding astrocyte, the average Ca^2+^ signal will be faint. Examples of such spike-triggered averages are shown in Figure 7 (E) for three cells labeled as “#”, “*”, and “&” in Figure 7 (A) used to center the time windows. Here, Ca^2+^ spikes in cell “#” was time-locked to activation of a single other cell, activation of cell “*” lead to repeated activation of several other cells with a stable delay, while activation of cell “&” did not repeatedly lead to activation of other cells.

Inspired by this cell-wide “repeatability” measure based on the contrast of spike-triggered averages, we defined a similar score for more a detailed mapping of whether activation in some region repeatedly lead to activation in other areas with stable time lags. To this end, we split the spatial domain into overlapping square windows of size 5 × 5 *µ*m and extracted Ca^2+^ dynamics from these patches. We then selected the patches where there were more than 10 Ca^2+^ spikes reaching at least 0.5 *µ*M [Ca^2+^]_*i*_ and created spike-triggered 50 s-long averages from the raster of Ca^2+^ signals in all patches. Percentage of all points with [Ca^2+^]_*i*_ *>* 0.2 *µ*M in such spike-triggered windows was used as the repeatability score for a given patch. The patches were then projected back onto the spatial template, with averaging of the values in overlapping areas between patches. This resulted in an automated mapping of areas leading to repeated downstream activation at the network level, revealing cells, serving as hubs in spreading multicellular events (Figure 7 F). As suggested in Brazhe et al (2018) for a simpler model, some spatial configurations of thick branches and leaflets can trigger persistent pacemaker-like activity, taking over the control of dynamics at the network level. We thus tested if alterations to the spatial template could lead to self-organization in a different spatial pattern of the centers of high repeatability (Figure 8). The cell labeled as “*” in Figure 7 was often activated from its direct neighbor to the right. Unlinking this cell from the network by setting to zero all contacts at domain boundaries (panel A) lead to a change in the repeatability map, decreasing the score for the cell “*” and increasing it for the three cells in the center. Unlinking the cell “*” from the network effectively silenced it, leaving only three cells with relatively high repeatability scores in the template.

**Figure 8.**
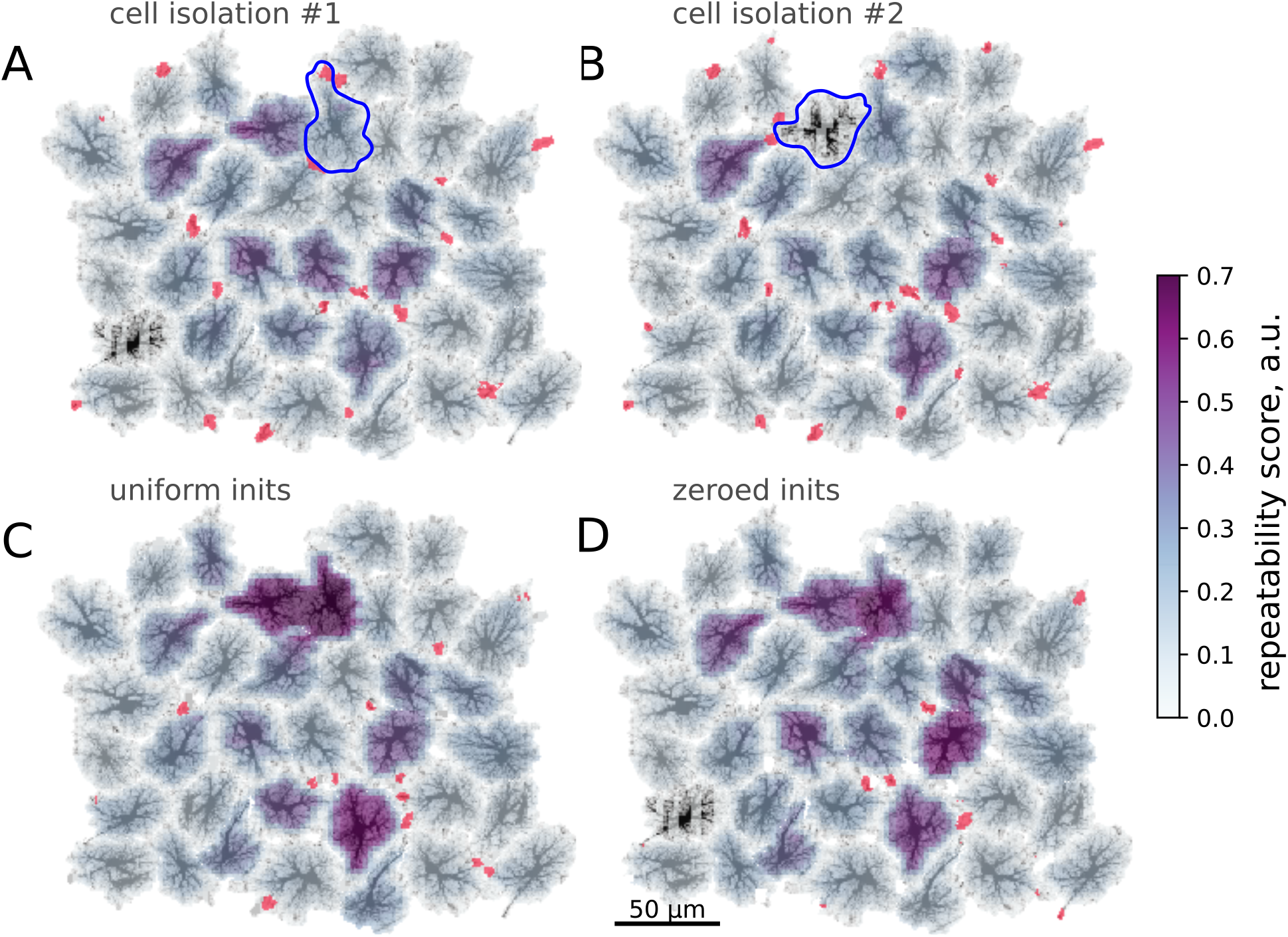
Modification of spatial templates changes activation patterns. Color-coding: repeatability score, red: wave initiation sites. (A–B) Unlinking single cells from the neighbors; Blue contours — isolated cells, red regions denote wave initiation sites as in Figure 4 (A). Isolation of the cell often activated the cell denoted “*” in Figure 7 lowers repeatability of the latter. (B) Isolation effectively silences the cell denoted “*” in Figure 7, which was a center of repeated patterns before. (C–D) Modifications of wave initiation sites. (C) Averaging AVF values within the activation sites increases repeatability of the waves, starting at the cells, isolated in (A–B) and one other cell. (D) Removing cell content from the spatial template within active initiation sites leads to a lower number of wave initiation sites and increased repeatability in several cells.

A subtler alteration of the spatial template can be directed at the sites of frequent Ca^2+^ transient initiation sites shown in red in Figure 8. Substituting the natural AVF profile at these sites with the average AVF value (panel C) or ablating cell content from these areas (panel D) lead to a dramatic reorganization of the Ca^2+^ initiation sites and the centers of high repeatability. In both cases the number of initiation sites was reduced, with some of the former sites being silenced and some new sites formed in the neighborhood of the previous sites. Also in both cases the inequality of repeatability score was increased, with a few cells showing very high values. We attribute this to the reduced number of initiation sites, leading to a more repeated activation sequences.

## 4 DISCUSSION

We proposed a spatially detailed model of astrocytic calcium activity, which reflects current understanding of the two distinct mechanisms of Ca^2+^ dynamics: excitable IP_3_-mediated exchange with ER in astrocyte soma and branches and plasma membrane exchange in the fine astrocytic processes and leaflets, sensitive to external conditions. Specifically, we suggest (i) an algorithm for data-based generation of 2D spatial templates matching realistic astrocyte morphology, and (ii) morphology-dependent spatially non-uniform parameter landscape for the calcium dynamics. To this end, we introduce the AVF parameter, which sets locally the relative input of the plasma membrane and ER based pathways and scales effective intracellular diffusion coefficients. The central idea underlying this separation is that astrocytes “sense” synaptic activity with fine processes, and it is where Ca^2+^ transients are relying on extracellular Ca^2+^ rather than intracellular stores, and where the bulk of IP_3_ can be produced, while thicker branches and somata provide the positive feedback gain mechanism for IP_3_-mediated Ca^2+^-induced Ca^2+^ release from ER. This mechanism separation is directly mapped to cell morphology in our approach.

We tested the suggested framework both at individual cell level and for algorithmically created multicellular astrocytic network templates. Our results show that the model is able to reproduce characteristic spatiotemporal patterns of Ca^2+^ dynamics driven by synaptic activity, represented by spatially uncorrelated point-sources of glutamate release coupled to focal Ca^2+^ entry, triggered by independent stochastic Poisson spike trains. At the single-cell level, the statistics of Ca^2+^ event durations and expansion areas turned out to have a power-law distribution resembling experimental data (Wu et al, 2014). Power-law statistics of the Ca^2+^ transients doesn’t directly follow from the model equations and is an emergent property of the interplay between astrocyte morphology and Ca^2+^ dynamics.

A notable feature of simulated Ca^2+^ dynamics in this system is spontaneously emerging stable patterns in both initiation and propagation of calcium transients, in tune to the co-active neuronal and astrocytic cells or repeating sequences of neuronal activation reported in slices (Sasaki et al, 2011, 2014). In agreement with Brazhe et al (2018) we observed morphology-dependent emergence of hotspots with persistent pacemaker-like activity, taking over control of the dynamics at larger scales. In single-cell templates these preferred initiation sites could lead to activation of either spatially confined microdomains or larger expanding Ca^2+^ areas, covering up to the whole cell. In multicellular systems we observed self-organized patterns of repeated calcium activity involving multiple cells. All cells in the template sharing the same equations and parameters, local differences in morphology of single astrocytes and geometry of astrocyte-to-astrocyte contacts favoured initiation of multicellular Ca^2+^ waves in some cells, followed by repeated sequences of cell activation as the Ca^2+^ wave sweeped across the network. Excluding some cells from the original template caused dramatic reshaping of reproducible activation patterns; removing or mashing up cell content in event initiation hotspots effectively reduced the number of active initiation sites and led to more stereotyped network activity. We conclude that with the same parameter set, the specific dynamical regime and role of an individual cell within astrocytic network is to a large extent defined by its morphology.

The presented model is a rather simplified representation of native astrocyte morphology and Ca^2+^ dynamics. It is less detailed, but also less computationally demanding than the framework proposed by Savtchenko et al (2018), allowing for simulations on a GPU-equipped laptop rather than on a supercomputer or a cloud. The major simplification of our model, dictating its limitations, is the reduction of real 3D astrocytic morphology to flat 2D spatial templates. The flattening was primarily done for the sake of computational tractability, but also conceptually matches single-plane imaging regime. The emergence of repeated activation patterns should not depend on the embedding space dimensionality, although the repeated propagation patterns can become more complex and elaborate in 3D; one can also expect different expansion rates and initiation probabilities for Ca^2+^ events in 3D. Notwithstanding, we argue that using 2D patterns can be a useful approximation. First, some astrocyte network systems can be regarded as effectively two-dimensional, e.g. astrocyte cultures or retinal astrocyte networks. Second, “true” astrocyte morphology can be regarded as less than three-dimensional, with the degrees of freedom limited by branching connectivity of the astrocytic processes, creating more or less independent astrocyte lobes and sub-domains. Another limitation comes from our approximation of the 3D mesh of fine astropil sponge by a continuous active medium, parameterized by astrocytic cytosol volume fraction. Effectively, we “glue” together the individual branches and leaflets, assuming that they are at least partly interconnected by autologous gap junctions and branch-to-branch loops. The idea of “loopy” or sponge-like organization of the astropil has experimental support (Wolff et al, 1998; Genoud et al, 2015; Arizono et al, 2020), and we therefore adopt the AVF framework to represent unresolved astrocyte processes, also accounting for the tortuosity of the sponge by AVF-dependent scaling diffusion coefficients.

Indeed, the mentioned simplifications are expected to limit the predictive power of the model with regard to event frequencies, scaling characteristics and propagation speed. On the other hand, the simulated patterns of single-cell calcium transients are qualitatively similar to that observed in a single focal plane, which suggests that this reduction seems to preserve main features of astrocyte dynamics, while it is worth to be investigated further at sub-cellular spatial scales in future work.

## Supporting information

Video1_network_template_creation

Video2_single_cell_dynamics

Video3_network_dynamics

## 5 ACKNOWLEDGMENTS

DEP and ARB acknowledge the support from the Russian Foundation for Basic Research, grant 19-515-55016, which covered the development of the modeling approach for spatial segmentation of inter-astrocyte functions. AYuV and ARB acknowledge support from Russian Science Foundation, grant 17-74-20089, which covered development of the numeric algorithm for creation of realistic spatial templates, biophysical interpretation of the model, and the collective astrocyte signaling analysis. DVV acknowledges support from the Russian Foundation for Basic Research, grant 16-32-50221 for mobility of young scientists, which covered model interpretation in terms of nonlinear dynamics, CUDA-based simulations of the model, and analysis of single-cell dynamics.

